# UNFOLDing Robustness, Plasticity, Evolvability and Canalisation of Biological Function

**DOI:** 10.1101/2025.03.03.641119

**Authors:** Debomita Chakraborty, Raghunathan Rengaswamy, Karthik Raman

## Abstract

A unique balance of seemingly contradictory properties like robustness and plasticity, or evolvability and functional canalisation, characterises biological systems. To understand the basis of these properties, we investigate gene regulation, which is at the core of biological function. We simulate dynamical models of over 190 million genetic circuits covering all possible three-gene circuit structures. Our computational pipeline classifies these circuits into functional clusters by matching their temporal response shapes. Thus, we generate a dataset linking circuit structure, parameters and a corresponding functional label. Our key finding is a finite list of 20 functions that three-node genetic circuits can perform. Moreover, the structure-parameter space for these circuits tends to be primed for responses that stabilise over time following a perturbation. Every structure exhibits potential for multifunctionality with a range of 2–17 functions contingent upon parameters. We quantify network degeneracy, showing that many structural changes can be made to circuits without altering function. We define three quantities: structural, parametric, and functional diversities. Using these diversities, we construct a UNified FramewOrk for reguLatory Dynamics (UNFOLD) to analyse four key biological properties—robustness, plasticity, evolvability, and functional canalisation. Using UNFOLD, we identify that only 6.5% of network structures are non-plastic, while parameter sets enabling parametric robustness exist for every three-node network. We identify functionally canalised circuits from structure pairs that can be interchanged for a large number of parameter sets without a change in function. Overall, our framework offers insights into the fundamental organisation of biological networks by thorough analysis of three-node networks.

**Significance Statement:** Biological systems exhibit remarkable properties like robustness, plasticity, evolvability, and canalisation. This study presents a unified computational framework to understand these properties by exhaustively exploring the design space of three-node genetic circuits, identifying that only 20 functions are achievable, and revealing a natural bias toward stability. We uncover key principles of network degeneracy and multifunctionality, highlighting the versatility of genetic circuits. By analysing structural, parametric, and functional diversities, we identify structural and parametric changes that can transition a genetic circuit from robust behaviour to plasticity or from being canalised to becoming evolvable. Our work advances theoretical insights into biological function. It provides a method to identify alternate designs and parametric conditions for genetic circuits, paving the way for the design of reliable synthetic genetic circuits.

## 1 Introduction

Biological systems demonstrate a fine balance of robustness, plasticity, evolvability and functional canalisation. Robustness is the ability of the system to maintain a function under changes in the structure and/or parameters of the system or environmental changes. Over two decades back, Wagner [1] suggested the possible bases of robustness in biological systems to be network structure and variability rather than redundancy introduced by duplicate genes. He elaborated on the coexistence of robustness and evolvability, citing that mutations can unleash evolutionary innovations by inching towards novel phenotypes even when they do not affect a particular phenotype immediately due to a neutral mutational space that confers mutational robustness to biological systems. The need for a unified framework with a strong mathematical basis to analyse biological robustness in all possible forms has long been recognised [2]. Evolvability may be defined as the potential to evolve or exhibit diverse phenotypes at the organism or population level. A comprehensive study of evolvability can be found elsewhere [3], along with methods to quantify this property from an evolutionary biology perspective. Plasticity is the property that allows organisms to tailor responses to internal or external cues dynamically. Waddington [4] coined the term canalisation to describe the low variance in phenotypes due to genetic and environmental variations in the wild-type, unlike strains of organisms manipulated in the laboratory.

In this work, we aim to construct a unified framework with a solid mathematical basis to analyse robustness, plasticity, evolvability, and canalisation at the level of gene regulation. For this, we consider a genetic circuit as a dynamical system whose response (“circuit function”) to a stimulus depends on the circuit structure and parameters [5–8]. We assume that mutations and epigenetic or environmental changes can affect gene regulation by changing the structure and/or parameters of genetic circuits, thereby influencing the circuit function, i.e. phenotype. In our unified framework, we do not directly quantify parametric robustness, plasticity, evolvability, and functional canalisation [9] but define conditions under which these properties are exhibited. These conditions are based on the changes in circuit structure and parameters. Our analysis does not refer to their underlying biological cause, which may be mutational, epigenetic, or environmental.

The space of all possible circuit structures and parameters constitutes the design space. We study the design space of all possible three-node genetic circuits based on the classic framework proposed by Tang and co-workers [10, 11], who studied three-node networks to identify the structural conditions necessary for perfect adaptation. Our focus, however, expands to exploring the global design space of three-node genetic circuits to find all possible functions. Most existing works in the literature aim to design or fine-tune a genetic circuit to achieve only one functionality of interest. One notable work that delves into the design space is the Design Space Toolbox V3, which adopts a systems-theoretic approach [12, 13]. Still, its assumptions regarding the kinetic equations may not accurately model underlying mechanisms and cannot be used to explore multiple stable states [14].

We develop a computational approach to explore a broader scope than the previous works. Firstly, we map the circuit structures and parameters with the circuit functions. We simulate the mathematical models of the circuits over millions of parameter sets and pass the simulated data through our computational pipeline for clustering circuits with similar temporal responses. The output of this pipeline is a dataset with circuit structure and parameters along with a functional label. Secondly, we use this labelled dataset to construct and use our unified framework for analysing the four biological properties– parametric robustness, plasticity, evolvability, and functional canalisation.

In Section 2, we describe our methodology for generating the simulated dataset and the computational pipeline we have used to process the dataset to get the labels. In Section 3, we discuss our key findings. Further details of our methodology and validation of our findings are provided in Supplementary Information.

## 2 Methodology

We develop a computational framework for studying the design space of genetic circuits. The main steps involved are generating time-course data by simulation and creating a computational pipeline for processing these data, as illustrated in Figure 1.

**Figure 1.**
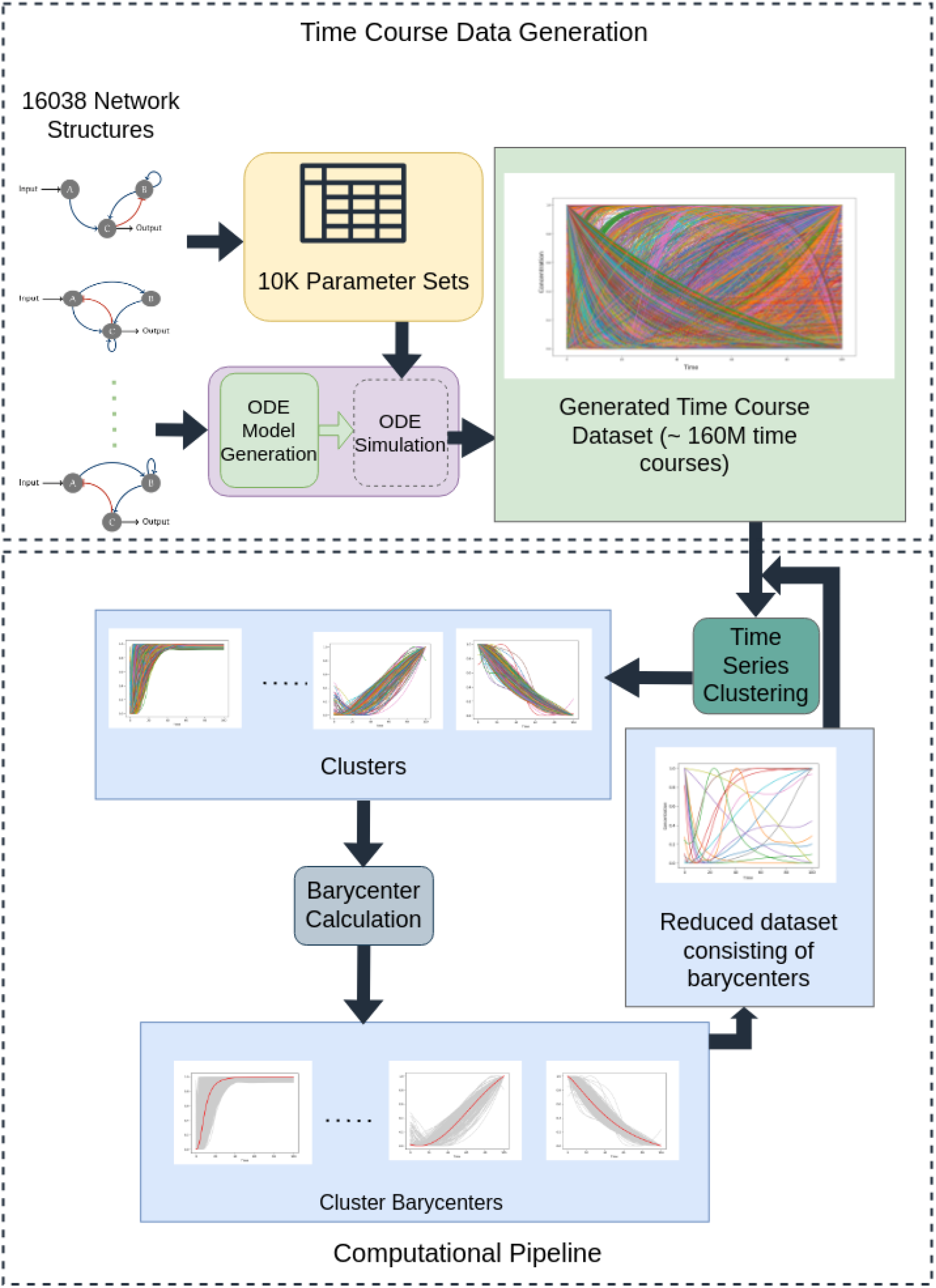
Workflow Diagram. We derive an ODE model for each of the 16038 networks from their adjacency matrices and simulate these models over three sets of 10,000 parameter sets each, sampled using Latin Hypercube Sampling. We pass the resulting time course dataset through a computational pipeline that performs time series clustering followed by barycenter calculation for each time course cluster. This constitutes the first iteration of the pipeline. (The barycenter corresponding to a cluster is shown in red.) For every subsequent iteration, the barycenters constitute a reduced dataset that serves as the input to the computational pipeline.

**Figure 2.**
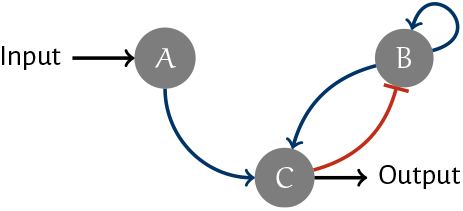
A representative three-node genetic circuit. *A, B, C* represent genes that interact by activation (shown as arrowheads) or inhibition (bar-heads). An activating input is applied to gene *A* while we study the expression of gene *C*.

### 2.1 Simulation for Time Course Data Generation

We represent a genetic circuit as a network with genes as the nodes. Each of the genes in the network expresses a transcription factor (TF) that either activates or inhibits the expression of the other genes in the network. These two types of interactions are represented by two types of edges in the network, *viz*., activation as arrowheads, and inhibition as bar heads. We label the three nodes as *A, B*, and *C*, where a step input is applied to node *A*, and hence, it is designated the input node, while node *C* is called the output node. The output gene expression over time for the given input, i.e. the expression of protein/TF from the output gene, *C*, is the focus of our analysis in this work. This approach is identical to that Ma *et al*. [10] proposed.

### 2.2 Setting up a Computational Pipeline

We simulate the ODE models to get each network’s concentration dynamics for 100 time points over the parameter sets for which a steady-state greater than 0.001 could be determined, as given in Supplementary Text Section S1.5. We construct a pipeline that recursively performs two operations: Time Series Clustering and Barycenter Calculation. We calculate the pairwise distances for time courses (min-max scaled) in our dataset using Dynamic Time Warping Distance (DTW-Distance) [15]. Subsequently, we perform K-Means clustering based on pairwise DTW distance. The number of clusters is determined by computationally locating the “elbow point” within a range of 2-100 clusters [16]. For each cluster identified by K-Means, the barycenter is the time series representative of the cluster. We calculate the barycenters of each of the clusters using Soft-DTW [17].

In the first iteration, we cluster the time courses obtained for each network over 10K parameter sets. Suppose a cluster with less than 10 time courses is found; we drop these time courses from further iterations as these represent network functionalities that cannot be achieved by even 0.1% of the parameter sets [10]. For all the remaining clusters for each network, the barycenters are calculated. The set of barycenters across all the networks then forms a reduced dataset for the second iteration of time series clustering. At the end of an iteration, we check if the shapes of all the barycenters are distinct, and if so, we terminate the execution at this iteration.

## 3 Results

We systematically examine all possible three-node networks by partitioning them into ten distinct partitions, each comprising 10% of them. The partitioned sets are obtained using a sampling procedure (detailed in Supplementary Text Section S1.2) that guarantees structural representativeness within each partition, effectively capturing the structural characteristics of the complete set of 16038 networks. We run two iterations of our computational pipeline with the time courses for each partition of networks. In the second iteration, the reduced dataset consists of about 15–16 clusters. We observe this across all the ten partitions of networks (Supplementary Text Section S1.6). Furthermore, we repeat the complete application of our computational pipeline to three sets of 10,000 parameter sets each to check for consistency in the results obtained (Supplementary Text Figure S1). Again, we find 15-16 distinct clusters over the ten partitions after the second iteration. Finally, we integrate the findings across the three runs to arrive at our consolidated results below.

### 3.1 Three-node networks perform 20 circuit functions

Our analysis identifies a set of 20 functionalities achievable with three-node circuits over 30,000 parameter sets, detailed in Supplementary Table S3. These represent distinct time courses, each embodying a unique shape that we interpret as distinct circuit functionalities illustrated in Fig. 3. A noteworthy observation is that 17 out of 20 circuit functions consistently emerge across all three pipeline runs, as detailed in Supplementary Text Section S1.6. Furthermore, we map each time course within a functional cluster to the specific network that produced it to establish a clear connection between the temporal dynamics and their underlying network structures.

**Figure 3.**
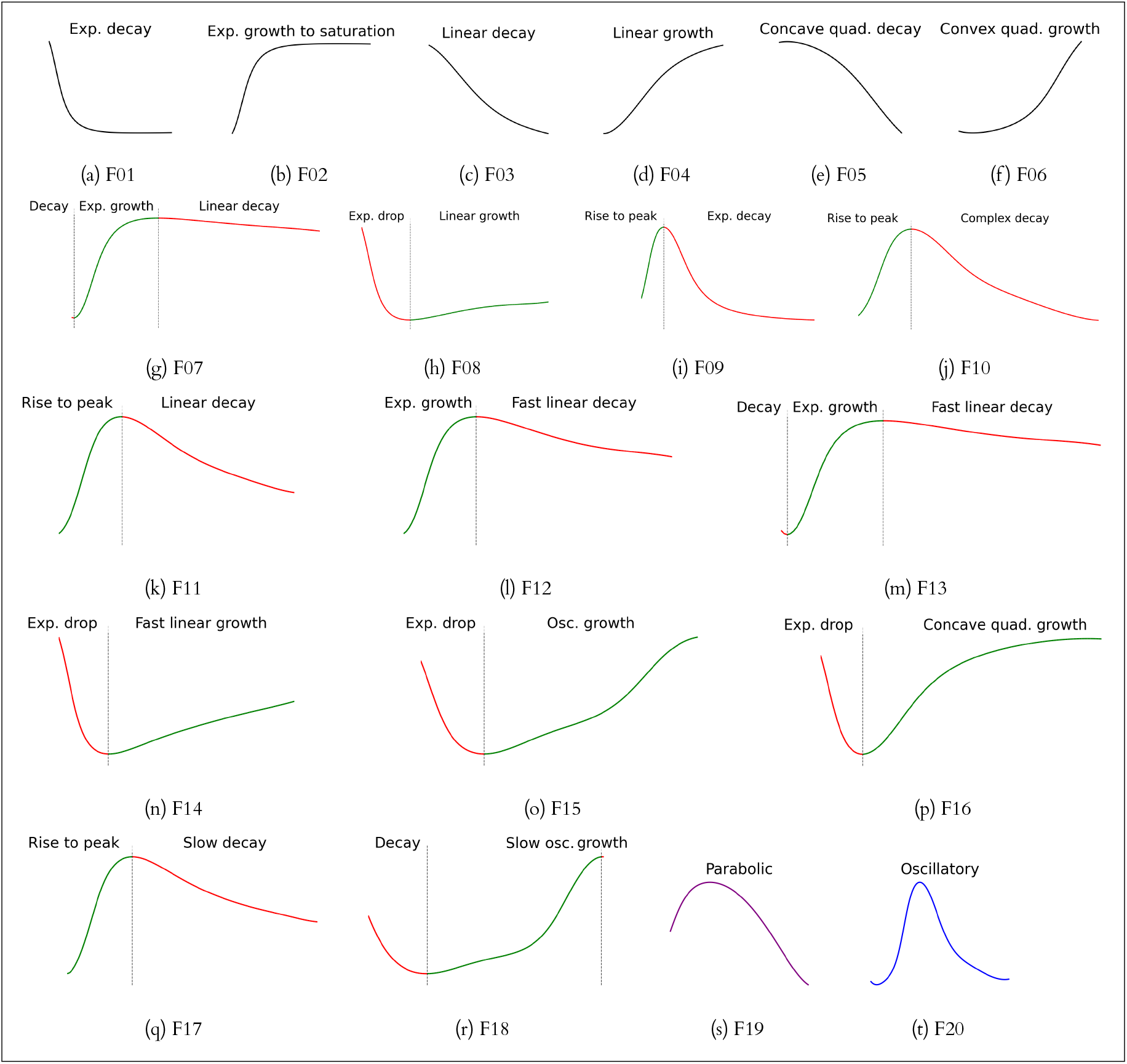
Three-node genetic circuit functions. The 20 distinct time courses characterise the functional space (in response to a step input) of all three-node genetic circuits. The description of each function is given in Supplementary Table S3, along with the percentage of circuits exhibiting the function.

### 3.2 The design space is primed to produce circuits that stabilise over time

Since circuit function depends on the network structure and the parameters governing the interactions within the network, we analyse the prevalence of the 20 functions exhibited by three-node networks in two ways: (a) in terms of network structure alone (network distribution) and (b) in terms of network structure with parameters defined (circuit distribution). In Supplementary Table S3, we show the high occurrence of circuits showing exponential decay (28.36%) and the circuits that saturate after exponential growth (27.37%) compared to the other circuit functions. This disparity suggests that the design space favours some functions over others and aligns intuitively with the observed tendency of biological systems to navigate toward stabilisation. The least frequent functions are characterised by oscillations (F20), a parabolic trajectory in time (F19), and decay followed by oscillating growth (F18). We note that the percentage of oscillatory networks represents only networks that are initially at a steady state and only respond to a step input with oscillations. Since we have eliminated any circuit that inherently shows instability (including oscillations) when we determine the initial conditions for our models, it may explain the small percentage of oscillatory networks. Moreover, we have eliminated any behaviour exhibited by a network for less than 10 out of 10000 parameter sets in the first iteration of our computational pipeline. These eliminated time courses may include oscillatory as well as adaptive behaviours. We separately analysed the simulated time courses to look for adaptive networks using criteria from existing works [10, 11]. The structural analysis of the adaptive networks reveals that using Hill kinetics, it is possible to achieve adaptation in three-node networks even without negative feedback loops or incoherent feedforward loops, unlike previously reported results by Shi [11]. The detailed findings on adaptive networks are given in Supplementary Text Section S1.7.

### 3.3 Every network is multifunctional

A notable observation in our analysis of three-node genetic circuits is that none of the networks displayed mono-functional behaviour. As we change the parameters of a given circuit, showing a particular function, it produces at least one new function at a different point in the parameter space. This finding highlights the intrinsic versatility of gene regulatory networks, showcasing their ability to exhibit diverse functionalities under varying parametric conditions. The number of distinct functions a three-node network performs spans an impressive range, from at least two to as many as 17 functions.

We further categorise the 20 circuit functions into five categories, *viz*., (i) monophasic, (ii) biphasic, (iii) triphasic, (iv) oscillatory, and (v) complex (Supplementary Table S4) based on the number of phases a time response can be divided into. The distribution of networks across the five categories is depicted in Supplementary Figure S2.1, while that of the circuits is shown in Supplementary Figure S2.2. The monophasic category encompasses networks that show either growth or decay. The biphasic category includes time responses that exhibit a period of growth (or decay) followed by a period of decay (or growth). The triphasic category includes responses with three phases of interleaved growth/decay. We consider oscillatory circuits a different category as they exhibit periodicity. We define a complex dynamic as one that shows a general trend of growth or decay but with oscillations or multiphasic behaviours superimposed on it.

Supplementary Figure S2 allows a nuanced analysis of the design space, revealing that while 5% of the networks manifest oscillatory behaviour, merely 0.02% of circuits exhibit this oscillatory behaviour. The percentage of circuits that exhibit the functions corresponding to each function category is much smaller than that of networks that fall in the corresponding category. This underscores the pivotal role of parameters in biological regulation determining a network’s function, complementing its structural characteristics. All 16038 networks are capable of exhibiting monophasic responses. However, networks that exclusively exhibit monophasic dynamics constitute only 21.8% of all networks. Interestingly, 53.03% of the networks can exhibit all the categories of functions under different parametric conditions except for oscillations, suggesting that an oscillatory response to a step input is a rare function. Other combinations of function categories are exhibited by <5% of the networks, with the rarest combinations being monophasic (I) and oscillatory (IV), or monophasic (I), biphasic (II) and oscillatory (IV). The rarest category of circuit function is oscillatory, while more than 90% of the circuits exhibit monophasic responses.

### 3.4 Network degeneracy allows a large number of structural changes with no change in function

We define network degeneracy as a measure of structural changes (edge addition/removal/change in the sign of an edge) that can be made, with no subsequent parameter adjustment, without causing loss of function. To quantify network degeneracy, we leverage our method of using a superset of parameter sets in 21 dimensions (for fully connected networks) to derive parameter sets for all 16038 networks (Supplementary Text Section S1.4). Removing an edge for a 21-dimensional parameter set makes the corresponding parameters zero, whereas adding an edge introduces the same parameter values from the 21-dimensional set under consideration. For simplicity, a change in the sign of an edge does not change the corresponding parameter values, i.e. the activation (inhibition) threshold and cooperativity become the inhibition (activation) threshold and cooperativity. For each parameter set in 21 dimensions, we track the number of structures that produce the same circuit function.

Supplementary Figure S3 shows the median numbers of structures over all parameter sets for each of the 20 functional clusters. While the network degeneracy depends on the function under consideration, we find a median of 60 structural changes across all functions.

### 3.5 A UNified FramewOrk for reguLatory Dynamics (UNFOLD)

We now describe a conceptual framework, UNFOLD, that unifies the analysis of the four properties of biological systems—parametric robustness, plasticity, evolvability and functional canalisation, using the results of our computational pipeline, as shown in Fig. 4. Considering a pair of circuits *C*_*i*_ and *C*_*j*_ with structures, parameter sets and functions given by (*N*_*i*_, *P*_*i*_, *f*_*i*_) and (*N*_*j*_, *P*_*j*_, *f*_*j*_), respectively, we define three quantities, *viz*., structural diversity (SD), parametric diversity (PD), and functional diversity (FD).

**Figure 4.**
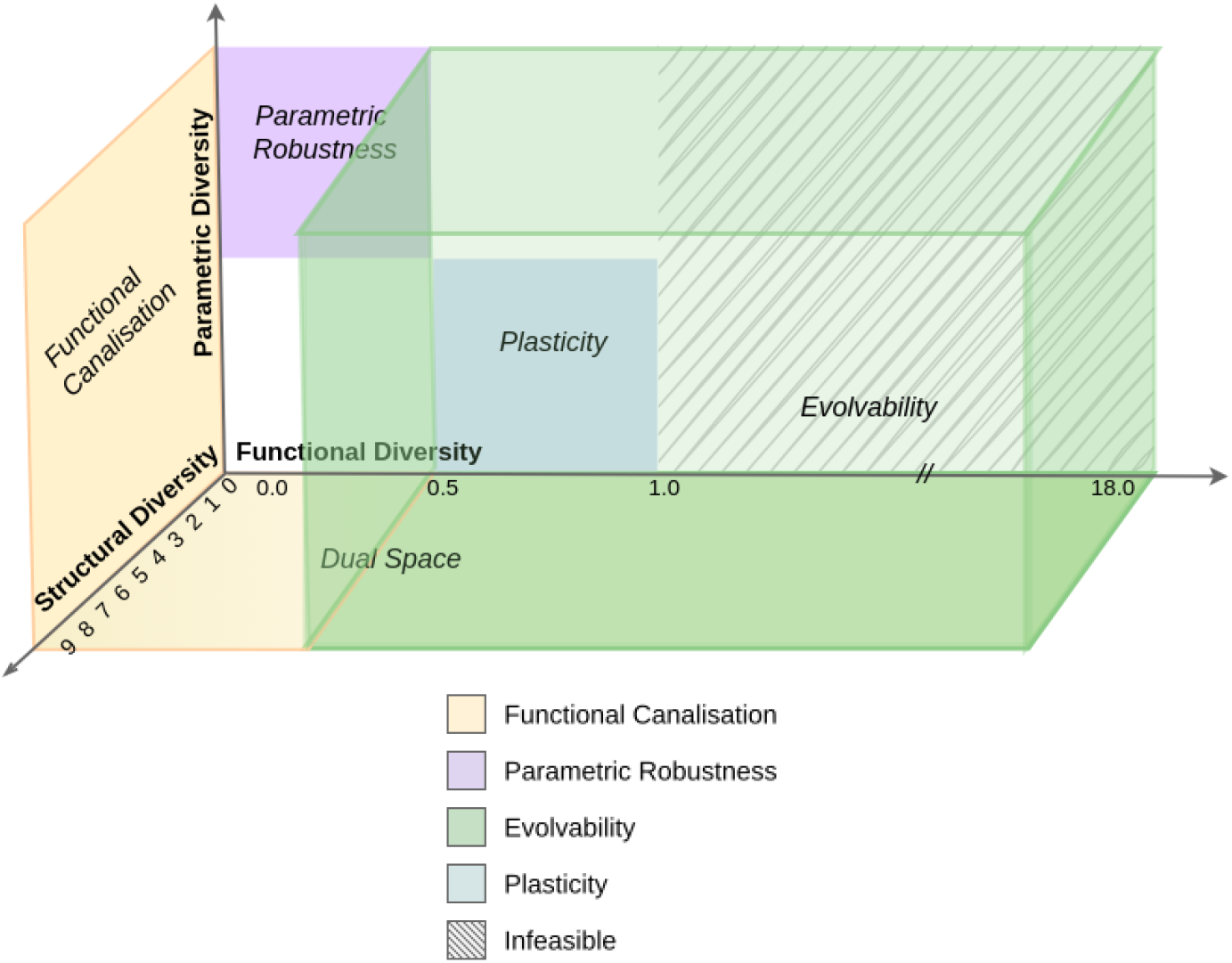
A UNfied FramewOrk for reguLatory Dynamics (UNFOLD). A conceptual framework for quantifying conditions for parametric robustness, plasticity, functional canalisation, and evolvability in terms of structural, parametric and functional diversities.

**Structural diversity** (*SD*_*ij*_) between *C*_*i*_ and *C*_*j*_ is quantified by the Hamming distance between the adjacency matrices *N*_*i*_ and *N*_*j*_ and assumes 10 discrete levels [0 *−*9].

**Parametric diversity** (*PD*_*ij*_) is quantified by the Euclidean distance between parameter sets *P*_*i*_ and *P*_*j*_ in the 21-dimensional parametric space.

**Functional diversity** (*FD*_*ij*_) is quantified using equation 1. We encode each network function as a *k*-hot binary vector of length 20 corresponding to the list of 20 possible functions with *k* indicating the number of functions exhibited by the network and ranging between 2 *−*17. We calculate the Hamming distance between the *k*-hot function vectors for *N*_*i*_ and *N*_*j*_ to obtain *HD*_*k−hot*_. Furthermore, we encode each circuit function as a *one*-hot binary vector of length 20 corresponding to the list of 20 possible functions. We define the Hamming distance between the function codes as *HD*_1*−hot*_, which is 0 if *f*_*i*_ and *f*_*j*_ are the same; else, 1.

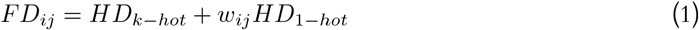

where *w*_*ij*_ is the weight quantifying the dissimilarity in functions depending on the function categories to which *f*_*i*_ and *f*_*j*_ belong, as given in Supplementary Table S5. FD is a discrete variable that assumes 37 unique values ranging between 0 *−* 18. For circuit pairs that share the same network structure (*SD*_*ij*_ = 0 since *N*_*i*_ = *N*_*j*_), *HD*_*k−hot*_ = 0, so FD can assume only 3 values *{*0.0, 0.5, 1.0}. This shows that constraining the structural diversity also constrains the functional diversity.

Using our computational pipeline, we have functionally labelled over 190 million three-node genetic circuits. Each data point in the conceptual space in Fig. 4 involves the calculation of structural, parametric and functional diversities for pairwise circuits. The number of computations involved in analysing all labelled circuit pairs is highly time-consuming and resource-intensive. We identify four properties of biological systems that can be analysed in terms of the three diversities: functional canalisation, parametric robustness, plasticity, and evolvability. These can be mapped to specific regions of the conceptual space in Fig. 4. Hence, we confine our analysis to these biologically relevant regions of this space and derive insights that will help us advance our understanding of the design space. The representation in Fig. 4 allows us to abstract the effect of mutations and/or epigenetic changes as changes in genetic circuit structure and/or parameters and enables us to dissect the role of structure and parameters in producing functional canalisation, parametric robustness, plasticity, and evolvability, in a unified framework.

#### 3.5.1 Conditions for parametric robustness and plasticity

Suppose mutations or epigenetic changes lead to changes in parameters but not the circuit structure; the circuits before and after the change constitute a pair with *SD*_*ij*_ = 0, *PD*_*ij*_ *≠* 0. For all such pairs of circuits, we calculate the parametric and functional diversity and find the overall mean parametric diversity *PD*_*overall*_*mean*_.

##### Parametric Robustness

We consider the circuit pairs for which *FD*_*ij*_ = 0, and *PD*_*ij*_ *> PD*_*overall*_*mean*_ to be robust as their functions/function categories remain unchanged for more than average changes in parameters.

##### Plasticity

Circuit pairs that have *FD*_*ij*_ *≠* 0 and *PD*_*ij*_ *< PD*_*overall*_*mean*_ are considered plastic as they exhibit diverse functions with relatively small changes in parameters.

We analysed the relationship between circuit structure and parametric robustness/plasticity in two ways for data points with zero structural diversity (SD). First, we identified the most frequently occurring network structures among robust and plastic circuit pairs. These predominant structures are illustrated in Supplementary Figures S4.1 and S4.3 for robust and plastic circuits, respectively. Second, we quantified the propensity of each network structure to generate robust or plastic behaviour. The most striking example, shown in Supplementary Figure S4.2, exhibits robustness in 52.67% of all circuit pairs sharing this structure. We found 209 distinct network structures maintaining robust behaviour across more than half of their parameter sets. In contrast, even the most plastic-prone structure exhibits parameter-dependent behaviour in only 40.75% of cases. Notably, our analysis revealed a fundamental asymmetry: while every three-node network structure can achieve robust behaviour with appropriate parameter selection, 1,043 structures (6.5% of the total) never display plastic behaviour, regardless of their parameters.

#### 3.5.2 Conditions for functional canalisation and evolvability

Suppose a circuit undergoes structural and parametric changes (*SD*_*ij*_ ≠ 0, *PD*_*ij*_ ≠ 0) due to mutations or epigenetic change but retains its function; then the circuit pair before and after the change represent a data point on the *FD* = 0 plane. To ensure *PD*_*ij*_ ≠ 0, we consider structural changes with at least one edge being added or removed (more details in Supplementary Text Section S1.8). A nuance to consider in this context is that when a mutation leads to a change in the structure of a circuit from *N*_*i*_ to *N*_*j*_, it inherently leads to changes in parameters corresponding to the edge changes while the parameters corresponding to the unaltered edges may or may not be affected. In our analysis, we do not distinguish between the two scenarios. Instead, we track only whether the circuit pair has zero functional diversity (*FD*_*ij*_ = 0).

#### Functional Canalisation

We interpret every point on the *FD* = 0 plane to be functionally canalised since they represent circuits that can maintain functionality even after changes in structure and parameters (Fig. 4). When a circuit pair having structures that share the same function vector (*HD*_*k−hot*_ = 0) exhibits the same function (*HD*_1*−hot*_ = 0), or it shows two distinct functions, but of the same category (*w*_*ij*_ = 0), the functional diversity is zero (Eq. 1). We find that canalised circuit pairs tend to have a peak SD between 4-6, indicating that a medium level of structural diversity is conducive to canalisation. Canalised circuit pairs can provide reliable alternate design options for achieving the same function in synthetic biology applications.

#### Evolvability

Circuit pairs for which *FD*_*ij*_ *≠* 0 and *SD*_*ij*_ *≠* 0 are defined as evolvable since they represent circuit pairs with both structural and parametric changes leading to new functions. The space representing evolvability with *FD ≠* 0 is shown in Fig 4.

The dual space encompasses the special case where *PD*_*ij*_ = 0. We discuss this space in detail in Supplementary Text Section S1.8.

## 4 Discussion

Our study presents the comprehensive exploration of the three-node genetic circuit design space, analysing over 190 million circuits to reveal fundamental principles of biological regulation. This unprecedented scale of analysis has led to discoveries that reshape our understanding of biological circuit design and function. While computational models have long enabled synthetic biological circuit design [18, 19], previous approaches like Tang and co-workers’ study of perfect adaptation [10] or Chiang’s KMFA pipeline focused on optimising circuit parameters for single functionalities [20]. Our work transcends the scope of these previous works by providing a complete map of the achievable functional space and establishing a unified framework for understanding circuit behaviour. The discovery that only 20 distinct functions are achievable by three-node genetic circuits represents a fundamental insight into the constraints of biological regulation. This finding, which emerged from analysing 30,000 parameter sets across all possible three-node network structures, suggests an inherent limit to the complexity achievable with three-node circuits. Remarkably, the distribution of these functions is highly non-uniform, with over 55% of circuits exhibiting either exponential decay or exponential growth to saturation. This strong bias toward stabilising responses to perturbation suggests an evolutionary preference for regulatory systems that can return to equilibrium over time. The implications of this finding extend beyond theoretical interest - it provides crucial guidance for synthetic biology efforts by defining the complete space of achievable functions with three-node circuits.

Our analysis revealed two fundamental properties of genetic circuits that have profound implications for both natural and synthetic systems. First, every network structure examined exhibits multifunctionality, capable of producing between 2 and 17 distinct functions depending on parameter configurations. This universal multifunctionality suggests that natural circuits may have evolved to exploit this inherent versatility, potentially switching functions through parameter changes rather than structural modifications. Second, we discovered significant network degeneracy, where multiple structural changes often preserve circuit function without requiring parameter adjustments. This finding has immediate practical applications in synthetic biology, offering multiple design options for achieving desired functions and suggesting strategies for engineering robust circuits. The unified framework we developed, based on structural, parametric, and functional diversity metrics, provides a computational approach to analysing fundamental biological properties. Previous studies by von Dassow [21] quantified the robustness of the segment polarity gene function through the count of random parameter samples that produce a phenotype. Wagner [22] suggested an antagonistic relationship between genetic robustness and evolvability but a positive correlation between phenotypic robustness and evolvability due to large neutral spaces indicating a many-to-one map between genotype and phenotype. However, Mayer showed using Boolean maps between genotype and phenotype that Wagner’s suggestion was valid only in special cases, while in general, phenotypes are more likely to have a trade-off between robustness and evolvability [23]. UNFOLD moved away from the genotype-phenotype paradigm to the circuit level to simultaneously consider structural, parametric, and functional aspects. This approach revealed that while every three-node network can achieve robust behaviour with appropriate parameters, 6.5% of structures are fundamentally non-plastic - a finding that is significant for synthetic biological applications where plasticity is not a desired property.

While well-characterised examples like the *lac* operon [24] and *kai* circadian genes [25] provide important functions arising from the interaction of three genes, our work reveals a much broader landscape of possible circuit behaviours. The practical relevance of our findings extends to critical areas like cancer research, as exemplified by the RAS gene family (KRAS, HRAS, and NRAS) [26, 27]. Our framework provides new ways to analyse such three-gene systems and identify potential therapeutic targets by understanding their regulatory dynamics within the complete functional space. Our approach complements experimental methods like scRNA-seq and ATAC-seq by providing a forward design perspective rather than inferring networks from data. Our comprehensive data repository serves as a valuable resource for synthetic biology, offering a complete map of structure-parameter-function relationships and enabling rational design of circuits with desired properties. This is particularly powerful when combined with our understanding of network degeneracy and alternative implementations, as it allows designers to choose optimal implementations based on practical constraints.

Several exciting avenues exist for expanding this work. While we focused on step input responses, future investigations could explore other input types and incorporate noise, which has been shown to produce richer dynamics [20]. The framework could be extended to larger networks, though the computational challenges would increase significantly. Most importantly, our findings suggest specific experiments to validate the predicted rare functional categories and test the practical implementation of canalised circuits. The key contribution of this study goes beyond providing a conceptual framework - it establishes fundamental limits on three-node circuit functionality, reveals universal properties like multifunctionality and network degeneracy, and offers practical guidelines for circuit design. By unifying the analysis of robustness, plasticity, evolvability, and canalisation, we bridge the gap between theoretical understanding and practical application, opening new avenues for basic research and synthetic biology applications.

## Code Sharing

All the code used for the current work is publicly available in the GitHub repository at https://github.com/RamanLab/UNFOLD-Framework.

## Acknowledgements

DC acknowledges the HTRA fellowship from the Ministry of Education, Government of India and Studentship from the Centre for Integrative Biology and Systems medicinE (IBSE), IIT Madras, India.

## References

[1] Wagner, A. (2005) Robustness and Evolvability in Living Systems. (Princeton University Press).

[2] Kitano, H. (2004) Biological robustness. Nature Reviews Genetics 5, 826–837.

[3] Hansen, T. F, Houle, D, Pavlicev, M, & Pélabon, C. (2023) Evolvability : a unifying concept in evolutionary biology? (The MIT Press).

[4] Waddington, C. (1957) The Strategy of the Genes. (Routledge), 1st edition.

[5] Raman, K. (2021) An Introduction to Computational Systems Biology: Systems-Level Modelling of Cellular Networks. (Chapman and Hall/CRC), 1 edition.

[6] Chakraborty, D, Rengaswamy, R, & Raman, K. (2022) Designing biological circuits: From principles to applications. ACS Synthetic Biology 11, 1377–1388.

[7] Brophy, J.A.N & Voigt, C. A. (2014) Principles of genetic circuit design. Nature Methods 11, 508–520.

[8] Slusarczyk, A. L, Lin, A, & Weiss, R. (2012) Foundations for the design and implementation of synthetic genetic circuits. Nature Reviews Genetics 13, 406–420.

[9] Kaneko, K. (2009) Relationship among phenotypic plasticity, phenotypic fluctuations, robustness, and evolvability; Waddington’s legacy revisited under the spirit of Einstein. Journal of Biosciences 34, 529–542.

[10] Ma, W, Trusina, A, El-Samad, H, Lim, W. A, & Tang, C. (2009) Defining network topologies that can achieve biochemical adaptation. Cell 138, 760–773.

[11] Shi, W, Ma, W, Xiong, L, Zhang, M, & Tang, C. (2017) Adaptation with transcriptional regulation. Scientific Reports 7, 1–11.

[12] Lomnitz, J.G & Savageau, M. A. (2016) Design space toolbox v2: Automated software enabling a novel phenotype-centric modeling strategy for natural and synthetic biological systems. Frontiers in Genetics 7, 118.

[13] Valderrama-Gomez, M. A, Lomnitz, J. G, Fasani, R. A, & Savageau, M. A. (2020) Mechanistic modeling of biochemical systems without a priori parameter values using the Design Space Toolbox v3.0. iScience 23, 101200.

[14] Adler, M, Szekely, P, Mayo, A, & Alon, U. (2017) Optimal regulatory circuit topologies for foldchange detection. Cell Systems 4, 171–181.e8.

[15] Meert, W, Hendrickx, K, Craenendonck, T, Robberechts, P, Blockeel, H, & Davis, J. (2022) DTAIDistance (Zenodo). 10.5281/zenodo.7158824.

[16] Arvai, K. (2020) kneed (v0.8.2) (Zenodo). https://github.com/arvkevi/kneed.

[17] Tavenard, R, Faouzi, J, Vandewiele, G, Divo, F, Androz, G, Holtz, C, Payne, M, Yurchak, R, Rußwurm, M, Kolar, K, & Woods, E. (2020) Tslearn, a machine learning toolkit for time series data. Journal of Machine Learning Research 21, 1–6.

[18] Elowitz, M.B & Leibler, S. (2000) A synthetic oscillatory network of transcriptional regulators. Nature 403, 335–338.

[19] Hasty, J, McMillen, D, & Collins, J. J. (2002) Engineered gene circuits. Nature 420, 224–230.

[20] Chiang, A.W & Hwang, M. J. (2013) A computational pipeline for identifying kinetic motifs to aid in the design and improvement of synthetic gene circuits. BMC Bioinformatics 14(Suppl 16), S5.

[21] von Dassow, G, Meir, E, Munro, E. M, & Odell, G. M. (2000) The segment polarity network is a robust developmental module. Nature 406, 188–192.

[22] Wagner, A. (2008) Robustness and evolvability: A paradox resolved. Proceedings of the Royal Society B: Biological Sciences 275, 91–100.

[23] Mayer, C & Hansen, T. F. (2017) Evolvability and robustness: A paradox restored. Journal of Theoretical Biology 430, 78–85.

[24] Jacob, F & Monod, J. (1961) Genetic regulatory mechanisms in the synthesis of proteins. Journal of Molecular Biology 3, 318–356.

[25] Ishiura, M, Kutsuna, S, Aoki, S, Iwasaki, H, Andersson, C. R, Tanabe, A, Golden, S. S, Johnson, C. H, & Kondo, T. (1998) Expression of a clock gene cluster kaiABC as a circadian feedback process in cyanobacteria. Science 281, 1519–1523.

[26] Huang, L, Guo, Z, Wang, F, & Fu, L. (2021) KRAS mutation: from undruggable to druggable in cancer. Signal Transduction and Targeted Therapy 6, 386.

[27] Simanshu, D. K, Nissley, D. V, & McCormick, F. (2017) RAS proteins and their regulators in human disease. Cell 170, 17–33.

