## Supplementary Information for "UNFOLDing Robustness, Plasticity, Evolvability and Canalisation of Biological Function"

### A Unified Framework to Dissect Robustness, Plasticity, Evolvability and Canalisation of Biological Function

19th March 2025

#### S1 Supplementary Text

##### S1.1 Network Structure Generation

A three-node network is represented by a $[3 \times 3]$ adjacency matrix having three possible elements $\{-1, 0, 1\}$, representing inhibition, no interaction, and activation, respectively. Thus, the structure of all possible three-node networks can be obtained by generating all possible $[3 \times 3]$ adjacency matrices with $\{-1, 0, 1\}$ as the only possible elements. There are $3^9 = 19683$ such adjacency matrices/networks. Of the 19683 networks possible, 3645 networks have no direct or indirect connection going from the input to the output node. Our analysis is done on the remaining 16038 networks [1, 2].

##### S1.2 Sampling Networks to Obtain 10 Partitions

We devise a strategy to sample ten disjoint subsets of the 16038 networks, each sampled set having 10% of the complete set of networks. To guarantee that the sampled networks represent the structures of all possible three-node networks, we ensure that the edge distribution in each of the ten sampled sets follows the edge distribution of the complete set of networks. One advantage of sampling the networks using this strategy is that it allows checking for recurring patterns in the temporal responses of these sets of networks.
Since each adjacency matrix element represents an edge in the network, the probabilities of an edge representing activation/inhibition/no interaction are obtained by the frequencies of the elements {-1, 1, 0} in the 16038 adjacency matrices as given in Table S1.

| Edge Probabilities | | | | | | | | | |
|---|---|---|---|---|---|---|---|---|---|
| Category | AA | AB | AC | BA | BB | BC | CA | CB | CC |
| -1 | 0.333 | 0.363 | 0.409 | 0.333 | 0.333 | 0.363 | 0.333 | 0.333 | 0.333 |
| 0 | 0.333 | 0.272 | 0.181 | 0.333 | 0.333 | 0.272 | 0.333 | 0.333 | 0.333 |
| 1 | 0.333 | 0.363 | 0.409 | 0.333 | 0.333 | 0.363 | 0.333 | 0.333 | 0.333 |

Table S1: **Edge probabilities table.** Probabilities of different categories of edges occurring in the complete set of 16038 networks.

We obtain a set of 1604 networks ( 10%) by weighted discrete random sampling of the nine adjacency matrix elements from the set {-1, 0, 1} where the weights are the edge probabilities given in Table S1. We note that the adjacency matrix elements are categorical variables and cannot be assumed to be normally distributed. So, we perform a Chi-square test for goodness-of-fit with the edge probabilities in Table S1 as the expected data and the edge probabilities of the sampled networks as the observed data. The $p$-value obtained for each edge is then found to be $\geq 0.98$. Thus, we fail to reject the null hypothesis, i.e., the sampled 10% networks have the same edge distribution as the set of all three-node networks. We use the above method to sample ten disjoint sets of networks, 9 containing 1604 networks and one dataset containing 1602 networks.

#### S1.3 ODE Model Generation

We assume that Hill kinetics govern the interactions between the genes and the TFs, similar to the seminal study of Ma *et al.* [1] and the subsequent work by Shi *et al.* [2]. We generate the corresponding ODE models using the adjacency matrix for a network.

$$\frac{dx}{dt} = v_x \left( \prod \frac{A_i^n}{A_i^n + K_i^n} \cdot \prod \frac{K_j^n}{{I_j}^n + K_j^n} \right) - \frac{x}{\tau_x} \tag{1}$$

The general equation of a Hill kinetic model is shown in Eq. 1 where $A_i$ is the concentration of an activating gene product, $A_i \,\epsilon\, \{\mathrm{A, B, C, I}\}$. $I_j$ is the concentration of an inhibiting gene product, $I_j \,\epsilon\, \{\mathrm{A, B, C}\}$. The parameters involved in the equations are given below:

$v_x$: maximal gene expression level for gene product $x$
$n$: cooperativity or Hill coefficient
$K_i/K_j$: activation/inhibition thresholds for the $i$-th activator/$j$-th inhibitor
$\tau_x$: half-life of gene product $x$

The activating and inhibiting interactions change the concentration of the gene product $x$, where $x \,\epsilon\, \{\mathrm{A, B, C}\}$. As shown in Eq. 1, we assume an AND logic for the interactions, i.e., the gene $x$ is activated only when all its activators are expressed to a high concentration and all its inhibitors are at a low concentration [2].

#### S1.4 Parameter Sampling

The Eq. 1 is transformed to Eq. 2 by non-dimensionalisation [2, 3]. This equation has a maximum of 21 network parameters when a network has the maximum number of possible edges, i.e., 9 edges. We use Latin Hypercube Sampling (LHS) to obtain 10,000 parameter sets from the 21-dimensional parameter space. We repeat this sampling 3 times to get three sets of 10,000 parameter sets each. All the networks are simulated using the same super-set of 10,000 parameter sets. While simulating the ODE model of a network that does not have the maximum number of possible edges, the parameters corresponding to a missing edge are made zero. This is equivalent to sampling parameters separately for individual networks using LHS. To validate this, we follow below steps:

$$\frac{dx}{dt} = \frac{1}{\tau_x} \left( \prod \frac{A_i^n}{A_i^n + K_i^n} \cdot \prod \frac{K_j^n}{{I_j}^n + K_j^n} \right) - \frac{x}{\tau_x} \tag{2}$$

1. We sample $10,000$ parameters in three dimensions using LHS and change the $z$-coordinates of all the samples to zero to obtain two-dimensional samples.

2. We generate a new set of $10,000$ samples using LHS in two dimensions.

3. We then check if the above two sets of samples follow the same distribution by performing a two-sample Kolmogorov-Smirnov goodness-of-fit test.

The $p$-values corresponding to each coordinate is 1.0. Furthermore, we repeated this validation for one-dimensional samples obtained by making the $y$ and $z$-coordinates of $10,000$ three-dimensional samples zero. In this case, too, the $p$-value for the two-sample Kolmogorov-Smirnov goodness-of-fit test is 1.0. Thus, the null hypothesis cannot be rejected, indicating that the samples obtained in steps 1 and 2 above follow the same distribution. Thus, using the super-set of parameters by making the parameters corresponding to missing edges zero is equivalent to re-sampling in the lower dimensional parameter space using LHS.
An activating step input (I) that changes from $0.06 - 0.6$ at the time, $t = 0$, is used for simulation. This input activates the input gene $(A)$ with parameters $K_I = 0.4$ and $n_I = 1$ for all the networks simulated. Parameter ranges used for the simulation of ODE models are given below:
$K : 0.001 - 1$ (sampled using a logarithmic scale)
$n : 1 - 4$ (integral values sampled linearly)
$\tau : 1 - 100$ (sampled using a logarithmic scale)

#### S1.5 Initial Conditions Determination

The initial conditions are determined by finding the steady states of a network for each of the 30,000 parameter sets with input value = 0.06. We simulate the ODE models using only the parameter sets for which the steady state of a network exists and is greater than 0.001.

#### S1.6 Consistency of results

After two iterations of running our computational pipeline for 10% of the network, we consistently get 15-16 clusters for all 10 sampled sets of networks and also across the three versions of parameter sets. After all 10 sampled sets of networks are run through the computational pipeline, we collect the barycenters to form a combined dataset that we again run through our computational pipeline. After this step, we get 18 barycenters for all three versions of parameter sets. To find out if the 18 functions are similar or not, we calculate the pairwise DTW distance, and if this distance is below a threshold, the corresponding pair of barycenters are very similar in shape and represent the same function. Based on this DTW distance, we then find the union of the 72 barycenters. This results in 20 distinct barycenters denoting the 20 circuit functions that three-node genetic circuits can perform.

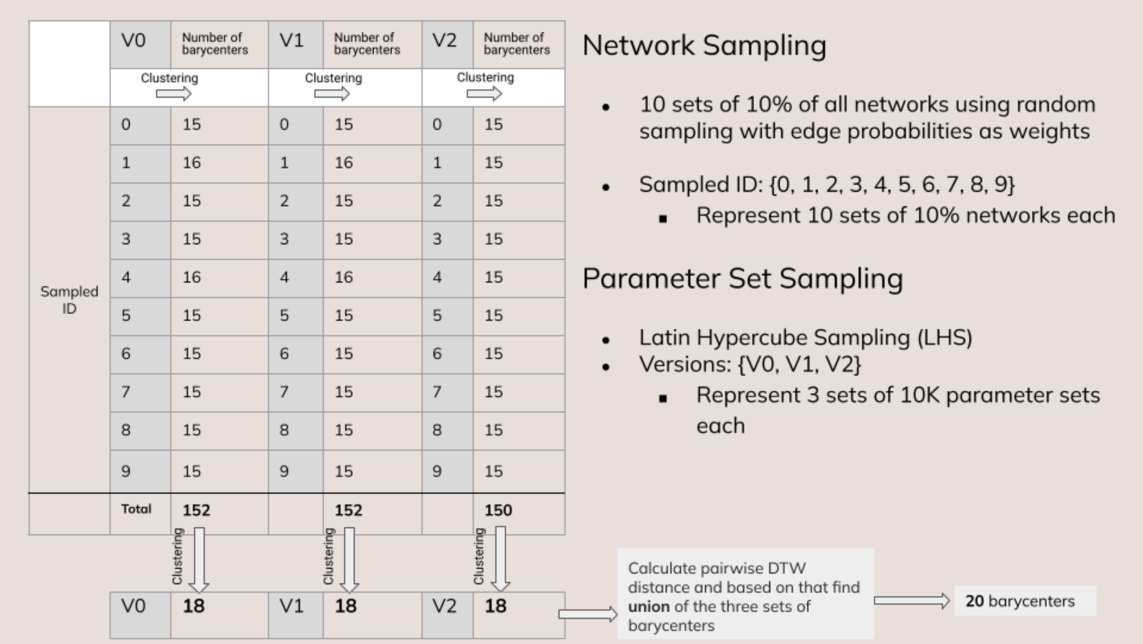


Figure S1: The 10 sampled sets of networks are labelled using the Sampled IDs from 0-9. The versions V0, V1, and V2 represent parameter sets. Each version contains 10,000 parameter sets.

#### S1.7 Analysis of Adaptation

For each of the 16038 networks, we filter out adaptive networks using the criteria of adaptation error ¡ 0.005 and response ¿ 0.2 as done by Shi and co-workers [2]. However, using these two criteria alone, we find oscillatory behaviours in the filtered time courses along with adaptation. So we further apply the filtering conditions used by Ma for detecting adaptation in enzyme regulatory networks [1]. These criteria filter out ill-behaved circuits, such as those which exhibit oscillations, or take a long time to stabilise (we choose this as more than 0.05% change even after 50% of the total simulation time). We then find the number of adaptive circuits for each of our three parameter sets (V0, V1, and V2). The results are listed in Table S2. We note that no network exhibits adaptation for more than 10 parameter sets. Our findings regarding adaptive networks are contrary to the results reported by Shi [2] as we find that networks with neither negative feedback loops (NFBL) nor incoherent feedforward loops (IFFL) are capable of exhibiting adaptation, in spite of the additional constraints we applied to remove ill-behaved circuits. We found 97 networks exhibiting adaptation across versions V0, V1, and V2, indicating that under different parametric conditions, a different set of networks are capable of exhibiting adaptation.

| Number of adaptive networks with | V0 | V1 | V2 |
|---|---|---|---|
| Negative Feedback Loops (NFBL) | 308 | 259 | 284 |
| Incoherent Feedforward Loops (IFFL) | 9 | 4 | 0 |
| Both NFBL and IFFL | 56 | 125 | 78 |
| Neither NFBL nor IFFL | 15 | 18 | 19 |
| **Total number of adaptive networks** | **388** | **406** | **381** |

Table S2: Adaptive networks having different structures across the three sets of parameters given by V0, V1, and V2.

#### S1.8 Dual Space

Given a circuit, when there is a change in the type of interaction from activation to inhibition or vice versa without introducing a new interaction or eliminating any existing interactions, we consider this a special case of structural change that produces *dual circuits.* If a circuit before and after such a structural change has at least one interaction type unchanged, then we term it a *partial dual* while a type change for all the edges results in a *complete dual.* The condition for duality is given below.

$$SD_{ij} \neq 0, \text{ and } PD_{ij} = 0 \tag{3}$$

In our sampling method, since we use parameter sets from a super-set, the numerical value of the parameters corresponding to the altered edges in duals do not change, i.e., although activation (inhibition) changes to inhibition (activation), the parameter values for these remain the same. $PD_{ij}$ being the Euclidean distance between a pair of parameter sets, for all dual pairs, $PD_{ij}$ is zero. If duals have zero $FD_{ij}$, they are functionally canalised duals, while duals with $FD_{ij} \neq 0$ fall in the evolvable category. Interestingly, functionally canalised duals produce the same function when one or more of its activating interactions is changed to an equal level of inhibiting interaction. When designing genetic circuits, understanding the dual space as defined in the proposed framework can help to identify design options for achieving a particular function, especially when the choice of some of the genes is constrained and only certain interactions can be manipulated. Notably, out of all possible ≈128 million pairs of 16038 three-node networks, we find 69174 pairs of networks that are complete duals. Moreover, we find 1343 parameter sets for which at least 10 (and a maximum of 44) complete duals exist for achieving the same function. Not all functions afford complete duals; however, for 16 out of the 20 possible functions, we could find at least one complete dual. We defer looking into the space of partial duals and constructing a library of alternate circuit designs using both complete and partial duals for the future. A particularly significant discovery is the existence of functionally canalised duals–circuits that maintain identical functions despite having opposite interaction types (activation versus inhibition). This finding has immediate practical applications in synthetic biology, offering multiple implementation strategies for desired functions and suggesting new approaches to circuit design.

### S2 Supplementary Tables

| Function ID | Function | % of circuits exhibiting the function |
|---|---|---|
| F01 | Exponential decay | 28.36% |
| F02 | Exponential growth to saturation | 27.37% |
| F03 | Linear decay | 17.57% |
| F04 | Linear growth | 11.40% |
| F05 | Quadratic decay - concave | 3.17% |
| F06 | Quadratic growth - convex | 2.43% |
| F07 | Decay followed by exponential growth followed by linear decay | 2.27% |
| F08 | Exponential drop followed by linear growth | 1.38% |
| F09 | Rise to peak followed by exponential decay | 1.04% |
| F10 | Rise to peak followed by complex decay | 1.03% |
| F11 | Rise to peak followed by linear decay | 0.87% |
| F12 | Exponential growth followed by fast linear decay | 0.82% |
| F13 | Decay followed by exponential growth followed by fast linear decay | 0.75% |
| F14 | Exponential drop followed by fast linear growth | 0.57% |
| F15 | Exponential drop followed by oscillating growth | 0.45% |
| F16 | Exponential drop followed by concave quadratic growth | 0.21% |
| F17 | Rise to peak followed by slow decay | 0.19% |
| F18 | Decay followed by slow oscillating growth | 0.05% |
| F19 | Parabolic | 0.05% |
| F20 | Oscillatory | 0.02% |

Table S3: **List of 20 functions that three-node genetic circuits exhibit.** The distribution of circuits over the design space is not uniform, with most functions exhibited by ¡2% of the circuits while the most exhibited functions are observed in ¿25% of the circuits.

| Category | Functional Cluster |
|---|---|
| I. Monophasic | F01, F02, F03, F04, F05, F06 |
| II. Biphasic | F08, F09, F11, F12, F14, F16, F17, F19 |
| III. Triphasic | F07, F13 |
| IV. Oscillatory | F20 |
| V. Complex | F10, F15, F18 |

Table S4: **Function Categories.** The 20 functions that three-node networks perform can be categorised based on the number of phases into which a given time response can be divided.

| $w_{ij}$ | **I.A** | **I.B** | **II** | **III** | **IV** | **V** |
|---|---|---|---|---|---|---|
| **I.A** | 0 | 0.5 | 1 | 1 | 1 | 1 |
| **I.B** | 0.5 | 0 | 1 | 1 | 1 | 1 |
| **II** | 1 | 1 | 0 | 1 | 1 | 1 |
| **III** | 1 | 1 | 1 | 0 | 1 | 1 |
| **IV** | 1 | 1 | 1 | 1 | 0 | 1 |
| **V** | 1 | 1 | 1 | 1 | 1 | 0 |

Table S5: **Weights table for calculation of functional diversity.** The rows and columns represent the categories as given in Table S4. We further sub-categorise the monophasic category (I) into I.A and I.B representing decay and growth, respectively. Sub-category I.A includes functions F01, F03, and F05 while I.B includes F02, F04, and F06.

#### S3 Supplementary Figures

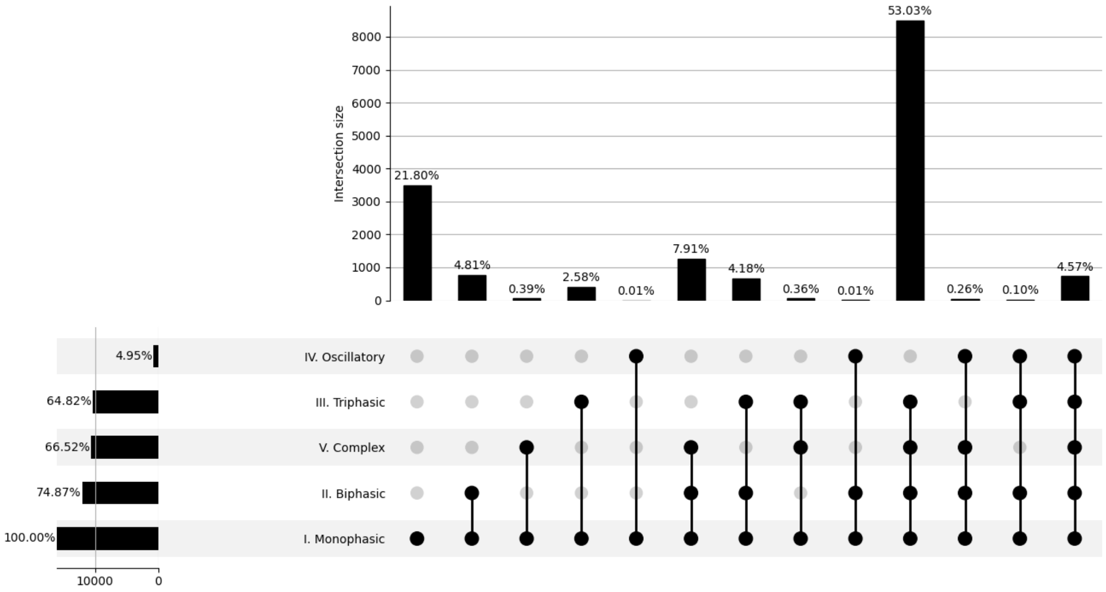


(S2.1) **Network distribution over function categories.**

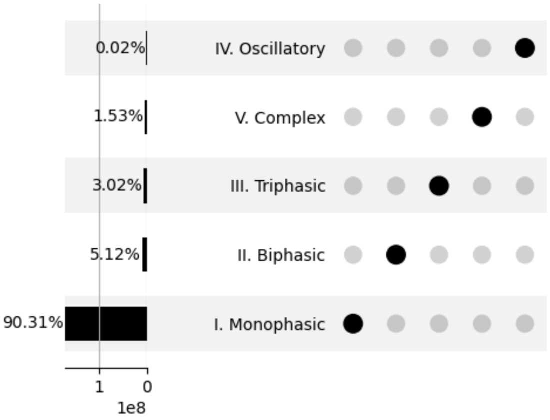


(S2.2) **Circuit distribution over function categories.**

Figure S2: Upset plots for network and circuit distributions over function categories.

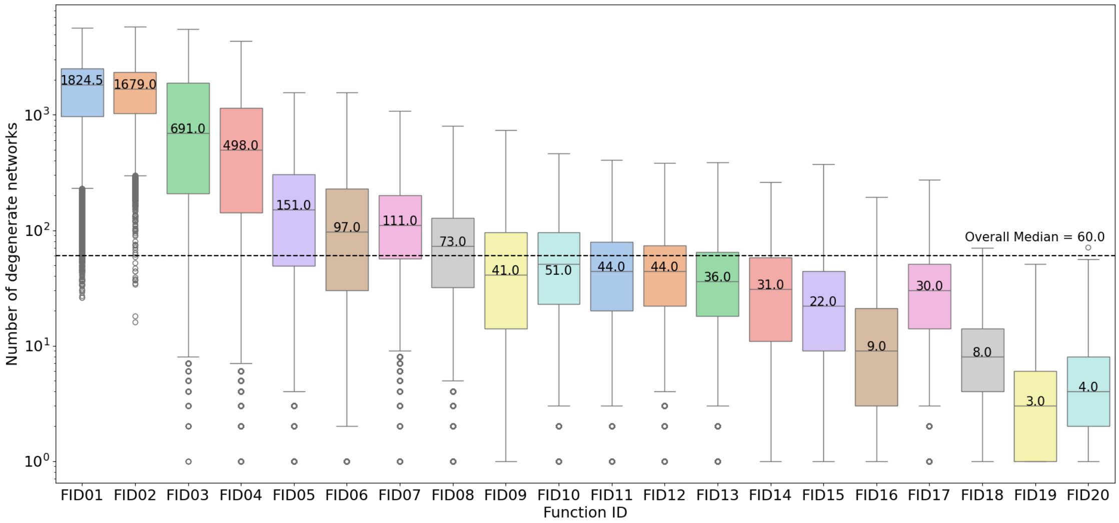


Figure S3: **Network degeneracy across different functions.** The median number of degenerate networks across all functions is 60, indicating that there can be 60 different structures with at least one addition/removal of an edge with/without change in the sign of edges (and parametric changes corresponding to only the added/removed edge, with other parameters remaining unchanged) without change in function. We find the median network degeneracy varies widely for different functions.

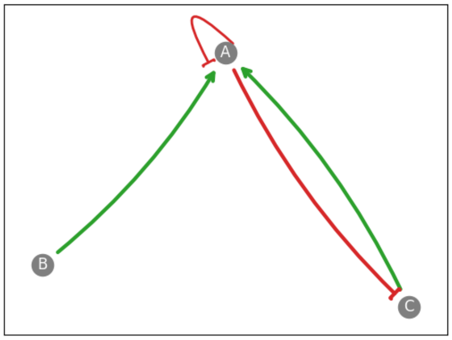


(S4.1)

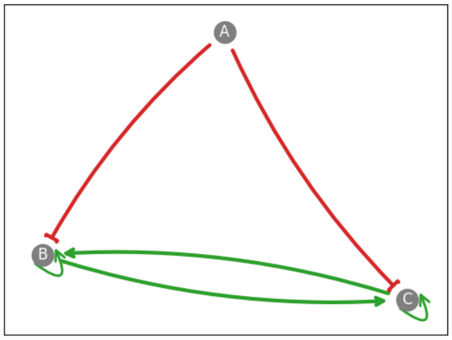


(S4.2)

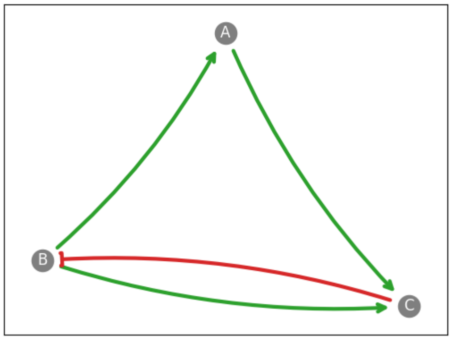


(S4.3)

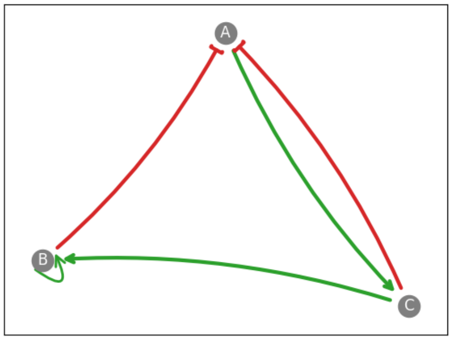


(S4.4)

Figure S4: **Structures of most robust and plastic circuits.**(1). The most common structure among all robust circuit pairs. (2). The structure that produces the highest percentage (52.67%) of robust circuit pairs out of all the circuit pairs that share this structure is shown in (b). The most common structure among all plastic circuit pairs is shown in (3). The structure that produces the highest percentage (40.75%) of plastic circuit pairs out of all the circuit pairs that share this structure is shown in (4).

### References


[1] W. Ma, A. Trusina, H. El-Samad, W. A. Lim, and C. Tang, "Defining network topologies that can achieve biochemical adaptation," *Cell*, vol. 138, pp. 760–773, 2009.

[2] W. Shi, W. Ma, L. Xiong, M. Zhang, and C. Tang, "Adaptation with transcriptional regulation," *Scientific Reports*, vol. 7, pp. 1–11, 2017.

[3] W. Ma, L. Lai, Q. Ouyang, and C. Tang, "Robustness and modular design of the drosophila segment polarity network," *Molecular Systems Biology*, vol. 2, pp. 1–9, 2006.
